# A unifying framework for quantifying and comparing n-dimensional hypervolumes

**DOI:** 10.1101/2020.11.21.392696

**Authors:** Muyang Lu, Kevin Winner, Walter Jetz

## Abstract

1. The quantification of Hutchison’s n-dimensional hypervolume has enabled substantial progress in community ecology, species niche analysis and beyond. While non-parametric methods for quantifying and comparing hypervolumes are popular, they do not support a partitioning of the different components and drivers of hypervolume variation. Here, we propose as alternative the use of multivariate normal distributions for the assessment and comparison of niche hypervolumes and introduce this as the multivariate-normal hypervolume (MVNH) framework.
2. The framework provides parametric measures of the size and dissimilarity of niche hypervolumes, each of which can be partitioned into biologically interpretable components. Specifically, We use 1) the determinant of the covariance matrix (i.e. the generalized variance) of a MVNH as a measure of total niche size, which can be partitioned into the components of univariate niche variances and a correlation component; and 2) the Bhattacharyya distance between two MVNHs as a measure of niche dissimilarity, which can be partitioned into the components of Mahalanobis distance between hypervolume centroids and the determinant ratio which measures hypervolume size difference.
3. We use empirical examples of community- and species-level analysis to demonstrate the new insights provided by these metrics. We show that the newly proposed framework enables us to quantify the relative contributions of different hypervolume components and to identify the drivers of functional diversity and environmental niche variation.
4. Our approach overcomes several operational and computational limitations of non-parametric methods and provides a framework that offers both unification and granularity in the assessment of niche volumes and differences, which has wide implications for understanding niche evolution, niche shifts and expansion during biotic invasions etc.

## Introduction

The n-dimensional hypervolume is one of the most fundamental (Hutchinson, 1957; Whittaker, Levin, & Root, 1973; Pulliam, 2000) and commonly used concepts (Díaz et al., 2015; Blonder, 2018; Pironon et al., 2018) in ecology and evolutionary biology. Hutchinson first proposed to describe species’ niche as an n-dimensional hypervolume in which a species can survive and reproduce (Hutchinson, 1957). The use of hypervolume was later extended to describe functional space and trait space (Lamanna et al., 2014; Laughlin, 2014; Pigot et al., 2020). The geometric features of n-dimensional hypervolumes (especially their size and dissimilarity) are associated with a wide range of hypotheses and applications in ecology and evolution. For example, the size of a climatic niche hypervolume is hypothesized to drive species diversification rates: in mammals, species with small niches (specialists) have been shown to have higher speciation rates and lower extinction rates than those with large niches (generalists) (Gómez-Rodríguez, Baselga, & Wiens, 2015; Rolland & Salamin, 2016). The size of climatic niche hypervolume is also hypothesized to drive the variation of geographic range size (Slatyer, Hirst, & Sexton, 2013; Saupe et al., 2015; Cardillo, Dinnage, & McAlister, 2019; Ficetola, Lunghi, & Manenti, 2020). Similarity between species’ environmental niche hypervolumes or functional trait hypervolumes is used to measure niche divergence or niche packing, which is hypothesized to determine species’ coexistence and species richness patterns (Serra-Varela et al., 2015; Pigot, Trisos, & Tobias, 2016; Castro-Insua, Gómez-Rodríguez, Wiens, & Baselga, 2018; Read et al., 2018). Niche similarity is also used for within-species comparisons such as assessing the impact of climate change (Tayleur et al., 2015; Gómez, Tenorio, Montoya, & Cadena, 2016; Zurell, Gallien, Graham, & Zimmermann, 2018) and whether there are niche shifts during biotic invasion (Lauzeral et al., 2011; Early & Sax, 2014; Guisan, Petitpierre, Broennimann, Daehler, & Kueffer, 2014; Davies, Hill, McGeoch, & Clusella-Trullas, 2019).

The quantification of niche volume and similarity has thus become an urgent pursuit for both theoretical investigation and real-world applications. Recently, non-parametric methods, such as kernel-density estimates (Blonder, Lamanna, Violle, & Enquist, 2014; Blonder et al., 2018; Eckrich, Albeke, Flaherty, Bowyer, & Ben-David, 2020; Mammola & Cardoso, 2020), dynamic range boxes (Junker, Kuppler, Bathke, Schreyer, & Trutschnig, 2016), minimum convex hulls, and support vector machines (Blonder et al., 2018), have been widely favored because of their minimal assumptions around the distribution of data and the existence of well-documented statistical packages supporting their exploration and use. In these methods, an n-dimensional distribution over environmental or trait space is typically converted to a hypervolume with a boundary defined by a particular isopleth (often 95%) of the distribution. Niche size is calculated as the volume of the enclosed space. Niche similarity of two hypervolumes can be quantified in multiple ways, either with the union and intersection of hypervolumes (Jaccard and Sorensen indices) or with the distances between the two hypervolumes (Loiseau et al., 2017; Mammola, 2019).

These non-parametric methods of quantifying hypervolumes have several methodological limitations: 1) niche volumes and similarity measures are sensitive to the parameterization of the underlying distribution and depend additionally on the choice of isopleth (e.g. 95% or 50%), 2) computational costs are high when the dimension of variables and sample size are large (Blonder et al., 2014), and most importantly 3) the non-parametric methods generally cannot say anything about the relative importance of the constituent components of the measures (but see Junker et al., 2016; Loiseau et al., 2017). For example, most non-parametric methods do not provide information on the impact of each individual dimension on the size of a hypervolume. For those that do (e.g. dynamic range box; Junker et al., 2016), the shape information (correlation among variables) on hypervolume size is ignored. Another example is the use of Jaccard/Sorensen indices and distance metrics for hypervolume similarity comparison. Though the two classes of metrics (Jaccard and Sorensen indices, Euclidean distances between niche centroids) are complementary in capturing hypervolume dissimilarity (Mammola, 2019), no analytical link exists between them for comparing their relative importance on overall hypervolume similarity. The Jaccard and Sorensen indices are used to calculate how much hypervolume is shared between the two hypervolumes, which are more informative when two hypervolumes intersect with each other. On the contrary, the Euclidean distance between hypervolume (centroid distance or minimum distance) is more informative when two hypervolumes are disjunct (Mammola, 2019).

The prevalence of non-parametric methods in quantifying hypervolumes also hampers the integration of empirical niche studies with niche theories because empirical metrics are disconnected from theoretical derivations. For example, in the theory of limiting similarity, the outcomes of competition exclusion and niche evolution are typically calculated based on competition coefficients derived from one-dimensional utilization curves (MacArthur & Levins, 1967; Leimar, Sasaki, Doebeli, & Dieckmann, 2013). However, the testable theoretical predictions derived by these studies rely on quantitative analyses in higher dimensions (Rappoldt & Hogeweg, 1980), meaning that with the prevalent non-parametric methods, questions such as “Does higher niche similarity among species lead to more competitive exclusion” could only be studied in a limited or qualitative way (D’Andrea & Ostling, 2016). Although various parametric dissimilarity metrics have been proposed (Morisita, 1959; Pianka, 1974; Lu, Smith, & Good, 1989), they were largely ignored because of their conceptual disconnection with the prevalent size metrics of hypervolume. Parametric measures that have clear analytical links to theory and which unify both size and dissimilarity measurements are urgently needed.

To fill in this gap, we propose a framework to quantify and partition niche volume and dissimilarity based on the assumption of multivariate normal (MVN) distribution of the niche variables under study and call this the MVN hypervolume or MVNH framework. Different to currently used non-parametric methods, this partitioning framework provides a powerful quantitative assessment of relative contributions of the constituent components in driving total niche variation. We chose the multivariate normal distribution because it is the most widely used assumption for niche assessment and modeling both in theoretical (MacArthur & Levins, 1967; Rappoldt & Hogeweg, 1980; Soberon & Nakamura, 2009; Jiménez, Soberón, Christen, & Soto, 2019) and empirical work (Swanson et al., 2015; La Sorte, Fink, & Johnston, 2018), which provides room for integration and synthesis of size and dissimilarity measures (Lu et al., 1989). We show that in the MVN hypervolume framework niche size is derived from the covariance matrix (Soberon & Nakamura, 2009), and that niche dissimilarity can be quantified from the Bhattacharyya distance (Bhattacharyya, 1946; Lu et al., 1989; Winner et al., 2018). We demonstrate through analytical derivation and empirical examples how the partitioning of the metrics reveals the key drivers of variations in functional diversity and environmental niche.

## Materials and Methods

In our MVNH framework, we propose that a multi-dimensional niche space be described by a multivariate normal distribution with probability density function:

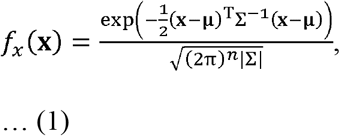

where **x** = {x_1_…, x_n_] is a vector in the n-dimensional environmental niche space, **μ** = {μ_1_ …, μ_n_] is the environmental niche centroid, Σ is the covariance matrix for the n-dimensional environmental niche, and |Σ| denotes the determinant of the covariance matrix.

### The size of a hypervolume

We define the size of a MVN hypervolume as the determinant of the covariance matrix |Σ|, also called the “generalized variance” (La Sorte et al., 2018). It has a geometric interpretation as the volume of the n-dimensional parallelepiped spanned by the column or row vectors of the covariance matrix. For example, the covariance matrix Σ and the generalized variance |Σ| of a two-dimensional MVNH are:

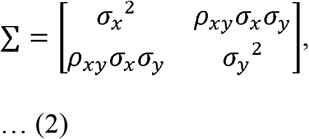

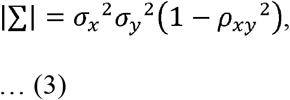

where 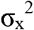 is the variance of variable x, 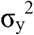 is the variance of variable y, and ρ_xy_ is the correlation coefficient between the two environmental variables x and y. When log-transformed, Equation 3 enables us to additively partition the size of a 2-dimensional hypervolume (in terms of the determinant) into three separate components (Fig. 1c): the variance of the first niche axis 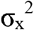, the variance of the second niche axis 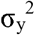, and a correlation component 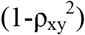. The correlation component could be seen as a shrinkage factor which shrinks the niche volume by a factor of 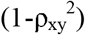. Thus, when two niche axes are independent, the niche size is just the product of the variances of individual axes 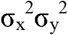. Similarly, for a three-dimensional hypervolume, the determinant of the covariance matrix is:

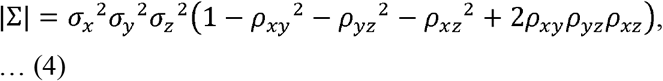

where is the correlation component. The factoring of the determinant generalizes easily to higher dimensions: the determinant is just the product of all individual univariate variances and a correlation component.

**Figure 1.**
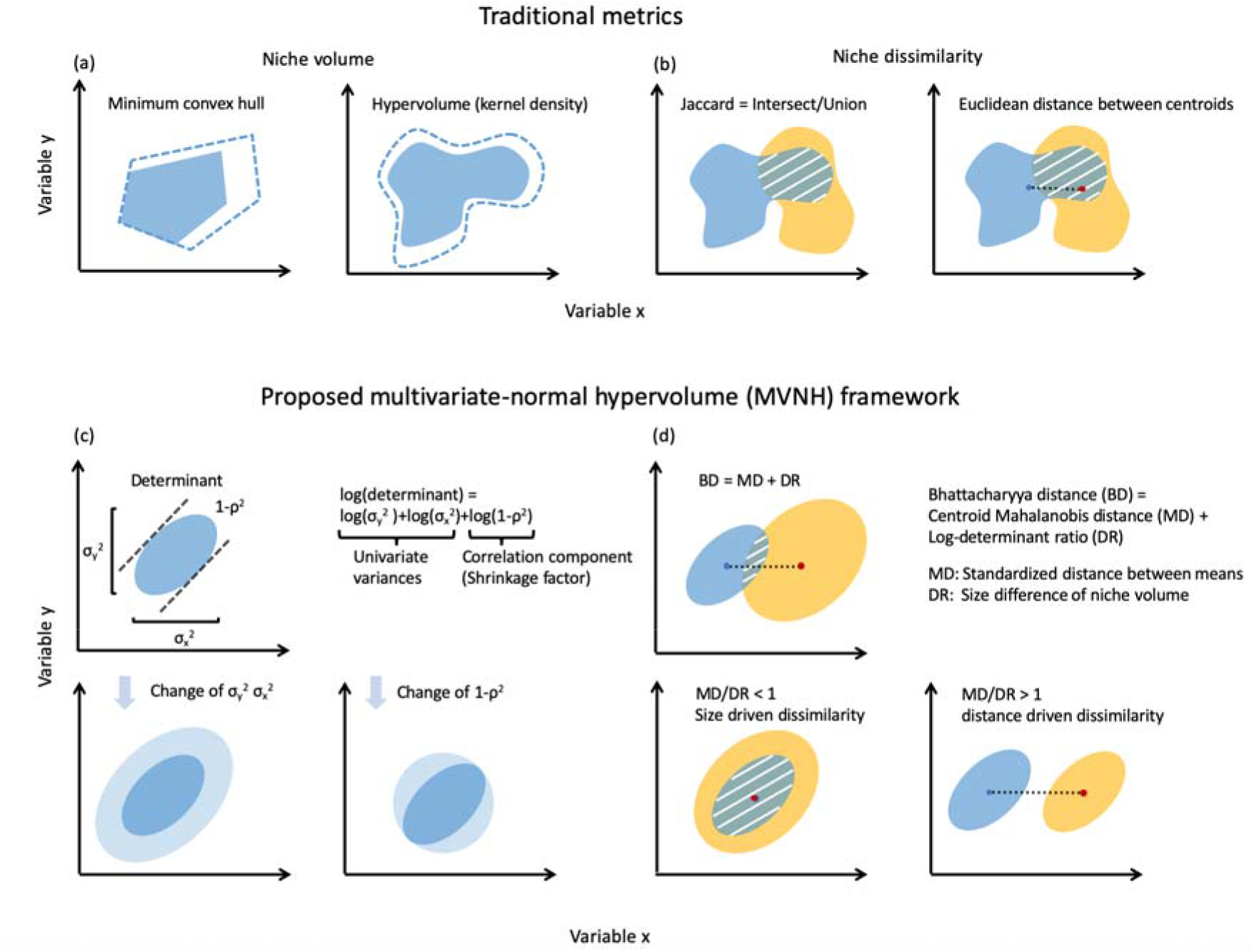
(a) Two commonly used non-parametric niche volume measures: the minimum convex hull and the kernel density estimated hypervolume. The filled area indicates 95% contour of the hypervolume while the dashed blue lines indicate 90% contour of the hypervolumes. (b) Two commonly used dissimilarity measures based on non-parametric volume estimates: Jaccard similarity and Euclidean distance between centroids. Orange and blue represent two hypervolumes. White hatches indicate intersecting area between two hypervolumes. (c) The volume of the 2-dimensional niche measured by the determinant of the covariance matrix of environmental variables, which is the product of three components: two univariate variances (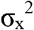 and 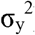) and a correlation component which has the effect of shrinking the volume by a factor of (1-ρ^2^). Light blue indicates change of hypervolume. (d) Niche dissimilarity of two hypervolumes measured by the Bhattacharyya distance, which is the sum of the Mahalanobis distance between the two niche centroids and a determinant ratio component measuring the differences of the niche volumes. The ratio MD/DR indicates the relative importance of centroid distance and size difference.

The determinant also has a close relationship with the standardized ellipse area (SEA, the ellipse with its major and minor axes defined by the principal components; Jackson, Inger, Parnell, & Bearhop, 2011): it scales with the SEA by a constant (π for two-dimensional niche and 4π/3 for three-dimensional niche). Factoring the determinant (as in Equations 3 and 4) can be used to assess the relative importance of the separate components in driving changes in the size of the hypervolume, such as when assessing whether individual trait axis or the constraint on the combination of traits is the major driver of community-wide trait diversity.

### The dissimilarity of hypervolumes

We suggest measuring the dissimilarity of two MVN hypervolumes with the Bhattacharyya distance (Bhattacharyya, 1946; Lu et al., 1989; Minami & Shimizu, 1999; Fieberg & Kochanny, 2005; Winner et al., 2018). The Bhattacharyya distance (BD) of two continuous probability distributions, *A* and *B* defined on the same domain 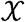 is:

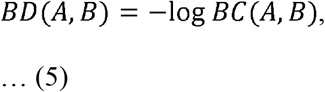

where BC is the Bhattacharyya coefficient:

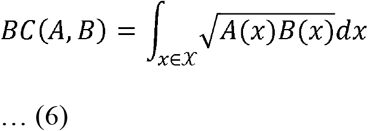

Therefore, for two probability distributions, the BC ranges from 0 to 1, and the BD ranges from 0 to ∞.

For two multivariate normal hypervolumes *A* and *B*, the BD has a closed form:

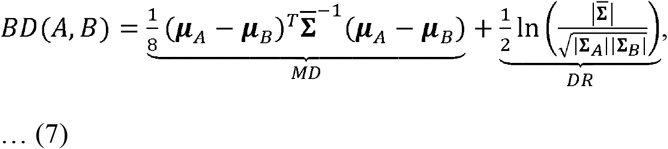

where μ_A_ and μ_B_ are the centroids of A and B respectively and 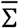 is the mean of the covariance matrices of A and B:

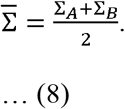

The first term in Equation 7 is the Mahalanobis distance (MD) between two hypervolume centroids, and the second term in Equation 7 is the determinant ratio (DR) which measures size difference between A and B (see Fig. 1d). If two hypervolumes have the same size, then the second term in Equation 7 will equal 0. As with Equations 3 and 4, factoring the Bhattacharyya distance according to Equation 7 can be used to assess the relative contributions of the Mahalanobis distance (centroid difference) and the determinant ratio (size difference) to the overall niche dissimilarity. We suggest that the ratio of Mahalanobis distance to determinant ratio (MD/DR), can be used as measure of the relative importance of centroid distance and size difference in driving overall dissimilarity.

It should also be noticed that the Mahalanobis distance component in Equations 7 could be linked to an existing partitioning framework which additively partitions the Mahalanobis distance into standardized Euclidean distances along PCA axes which measure the relative importance of each PCA axis (Calenge, Darmon, Basille, Loison, & Jullien, 2008; Mahony, Cannon, Wang, & Aitken, 2017).

### Estimators

We used sample means and sample covariances as estimates of μ_A_, μ_B_, ∑_A_, and Σ_B_ for the calculation of the determinant of the covariance matrices and the Bhattacharyya distance with its partitioned components. These plug-in estimates of the Bhattacharyya distance are known to be biased, but the bias is small with low dimensionality (less than 4) of data (Lu et al., 1989; Minami & Shimizu, 1999). For the purpose of demonstrating the utility of the partitioning framework, our empirical analysis proceeds without applying existing biascorrection methods which have been presented in the literature such as estimating the covariance matrix using Bayesian methods (Swanson et al., 2015), and bias-correction techniques for the Bhattacharyya distance (Minami & Shimizu, 1999; Winner et al., 2018). Calculation of both metrics only requires a few lines of codes in basic R, which is attached in the supplement (Appendix 1).

We also recognize other dissimilarity metrics based on multivariate normal distributions which share a similar form with the Bhattacharyya distance and could be partitioned into a Mahalanobis distance component and a determinant ratio component (see Lu et al., 1989), such as the MacArthur-Levins measure (MacArthur & Levins, 1967):

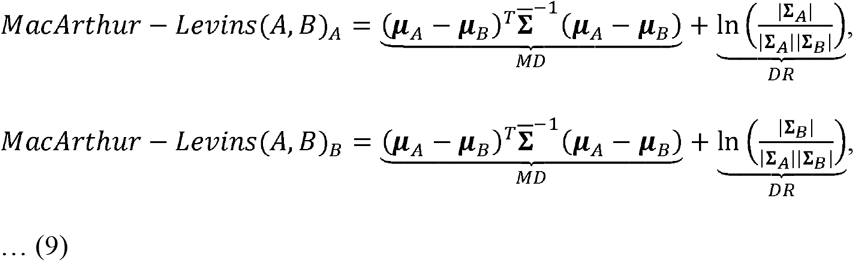

the Pianka’s measure (Pianka, 1974):

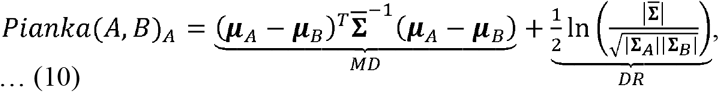

and the Morisita’s measure (Morisita, 1959):

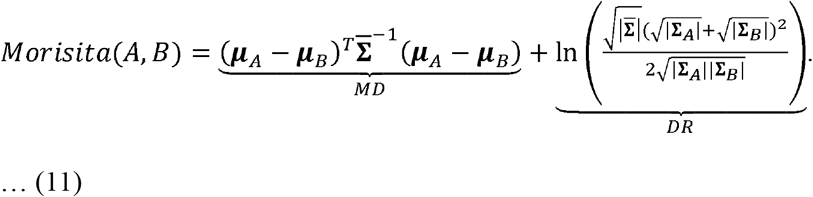

These metrics differ in their probabilistic interpretations, weighting of each component, and how they calculate the determinant ratio component. We will focus on the use of the Bhattacharyya distance in this paper because it was most studied with known statistical properties (Lu et al., 1989; Minami & Shimizu, 1999; Winner et al., 2018), and also has a straightforward interpretation as the integration of geometric means of two probability distributions (Equation 6).

### Empirical examples

We will use empirical examples of functional diversity and environmental niche to show the novel insights provided by the partitioning framework especially in quantifying the relative importance of univariate variance and correlation in driving size variation; the relative importance of centroid distance and size difference in driving hypervolume dissimilarity.

### Community example: Functional diversity and turnover

We used a trait dataset of 36 annual plant communities in Western Australia (Data availabile from the Dryad Digital Repository https://doi.org/10.5061/dryad.76kt8; Dwyer & Laughlin, 2018) to demonstrate how functional diversity (alpha diversity) and turnover (beta diversity) can be measured in the MVN hypervolume framework using the determinant and the Bhattacharyya distance, respectively. The three measured traits for each species were specific leaf area (SLA), maximum height (MH), and seed mass (SM). We then assessed each component of the trait volume (SLA variance, MH variance, SM variance and the correlation component) as predicted by growing season available moisture, mean growing season minimum temperature, and soil surface dispersion (dispersion is a measure of heterogeneity) in multiple linear regression. We used the coefficient of variation of each component to assess the relative contribution of each trait volume component to the variation of total trait volume, because the individual niche variances are scalable and not directly comparable with each other. We assessed the environmental drivers of the Bhattacharyya distance (functional turnover) components using multiple regression on distance matrices (MRM) based on a partial Mantel test (Lichstein, 2007).The regression coefficients of MRM are calculated with ordinary least square regression, but the p-values of the coefficients are calculated with matrix permutation tests. The predictor variables were calculated as the Euclidean distances for each environmental variable between sites.

### Species example: Environmental niche breadth and dissimilarity

We studied 260 bird species in the Western Hemisphere with occurrence data extracted from eBird (Sullivan et al., 2009), environmental layers obtained from CHELSA (monthly mean temperature and precipitations for breeding seasons, https://chelsa-climate.org/), and MODIS EVI (https://modis.gsfc.nasa.gov/data/dataprod/mod13.php) at 1km resolution to assess the drivers of realized environmental niche breadth (volume) and niche dissimilarity between species. 1000 random points were sampled and spatially thinned to 10km distance using the ‘spThin’ package (Aiello-Lammens, Boria, Radosavljevic, Vilela, & Anderson, 2015) in R for extraction of environmental data. We then investigated the drivers of environmental niche breadth components using ordinary multiple linear regression, and assessed the drivers of the Bhattacharyya distance components using MRM. We obtained body mass and dietary niche breadth data from the ‘EltonTrait’ dataset (Wilman et al., 2014). Area and centroid absolute latitude of geographic range were calculated from IUCN range maps (https://www.iucnredlist.org/resources/spatial-data-download).

### Comparisons to non-parametric metrics

For both the community and niche examples we compared the newly proposed size and dissimilarity metrics from the MVNH framework with measures based on the 95% isopleth of a box kernel density estimator using the ‘hypervolume’ package in R (Blonder et al., 2018). For the species environmental niche example, we also compared the determinant of the MVNH with the volume of minimum convex hull (Fig. 1a); for the community functional diversity example, we additionally compared the determinant with three commonly used functional diversity measures: functional divergence, functional dispersion, and functional evenness (Laliberté, Legendre, & Shipley, 2015). For both examples, we compared the Bhattacharyya distance with Jaccard and Sorensen similarities, centroid distances (Fig. 1b) and minimum distances between hypervolumes.

## Results

We use the MVNH framework to firstly analyze drivers of community-level functional diversity and functional turnover, and to secondly assess drivers of species-level environmental niche breadth and dissimilarity. Finally, we show the correlations between the proposed metrics with commonly used non-parametric measures.

### Community level: Functional diversity

We compared functional diversity (measured as the determinant of the trait covariance matrix) and functional turnover (measured as Bhattacharyya distance) among 36 plant communities.

For a two-community example (site 1 and site 2), we partitioned the functional diversity of each site into four components: the univariate variance of each of the three traits (specific leaf area, SLA; maximum height, MH; and seed mass, SM) and the correlation component following Equation 4. Component-wise differences between the two sites add up to the difference in total trait volume (the determinant of the environmental niche covariance matrix) on the log-scale (Fig. 2a).

**Figure 2.**
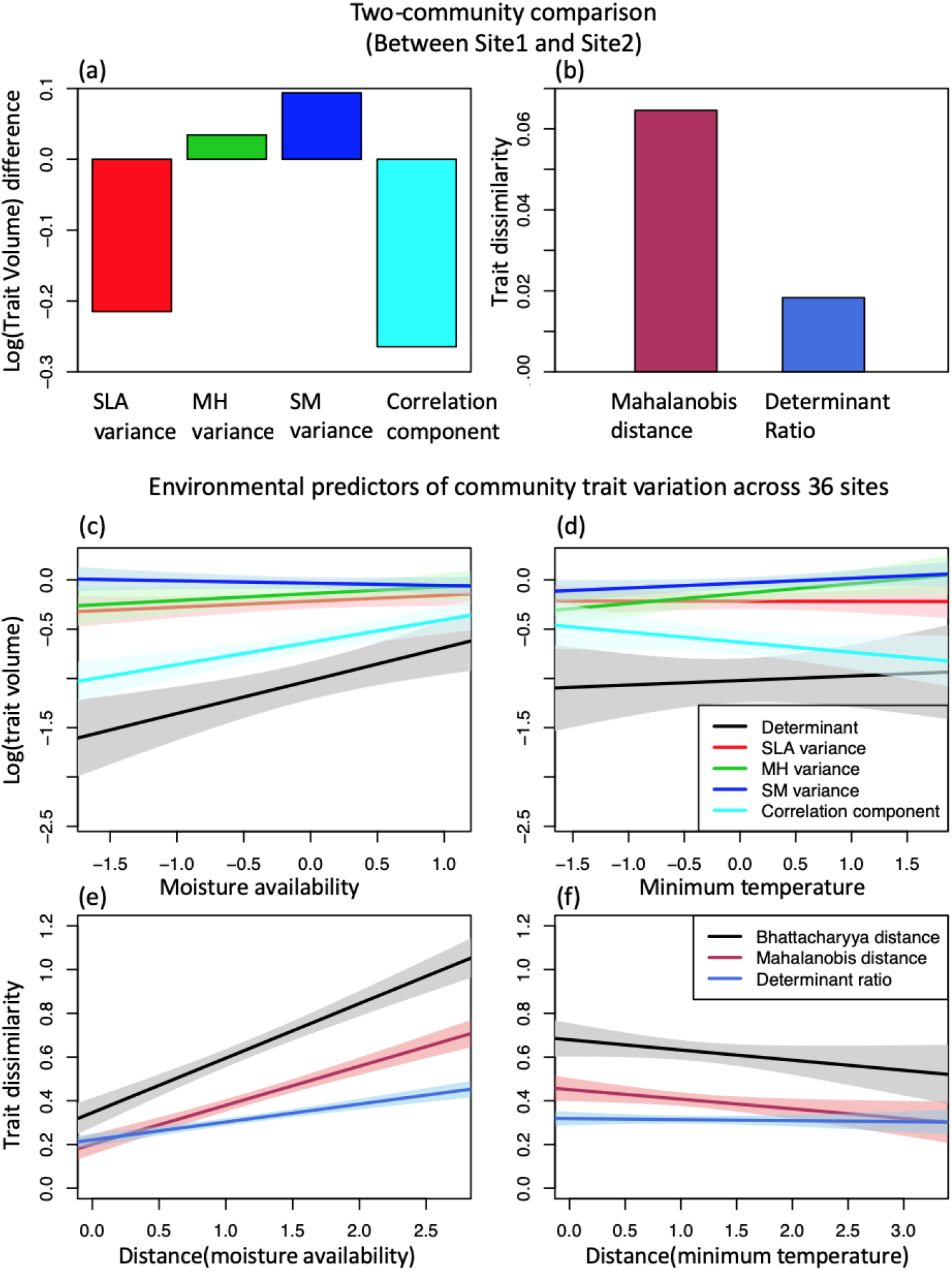
Community-level functional diversity assessment. (a) Component-wise differences between site 1 and site 2 in a) functional trait volume (e.g. functional diversity) and b) Bhattacharyya distance of functional traits. (c)-(f) Change of trait volume and pairwise community-level trait dissimilarity among all 36 sites of annual plant communities with environmental variables in southwest Western Australia. Solid lines are the fitted linear regression. Shaded area represents 95% confidence interval of linear regression. See table 1 for summary statistics.

For the 36-community assessment, the coefficient of variation for each trait volume component across the 36 sites was 0.31 for the correlation component, and varied between 0.28 (MH) and 0.16 (SM) for the individual trait variances. This suggests that the correlation component is the largest driver of the variation in functional diversity among sites. By relating individual trait variances and the correlation component to environmental predictors, we identify mean moisture availability and, less so, minimum temperature as key correlates of trait diversity variation (Fig. 2c, d). The variances of all three traits differ substantially in the strength and direction of their respective associations with environmental predictors. Linear regression shows that none of the examined environmental variables predict the variation of SLA variance (Table 1). Both minimum temperature and moisture availability are positively correlated with maximum height variance, whereas only soil surface dispersion is positively correlated with seed mass variance. Moisture availability is also positively correlated with the correlation component. The overall trait volume (the determinant) is only correlated with moisture availability (Table 1).

**Table 1.** Summary statistics for multiple linear regression of community-level trait volumes, and multiple regression on distance matrices on community-level trait dissimilarity. Specific leaf area, maximum height, seed mass variances, the correlation component and the determinant are log-transformed. Adjusted R^2^ is used. Only significant predictors are reported.

| Response | Predictor | Estimate | p-value | $R^2$ |
| --- | --- | --- | --- | --- |
| Specific leaf area variance | -- | -- | -- | -- |
| Maximum height variance | Minimum temperature | 0.125 | <0.01 | 0.19 |
|  | Moisture availability | 0.102 | <0.05 |  |
| Seed mass variance | Soil surface dispersion | 0.066 | <0.05 | 0.25 |
| Correlation component | Moisture availability | 0.228 | <0.001 | 0.37 |
| Determinant | Moisture availability | 0.369 | <0.001 | 0.27 |
| Mahalanobis distance | Distance (moisture availability) | 0.18 | <0.001 | 0.20 |
|  | Distance (minimum temperature) | -0.07 | <0.05 |  |
| Determinant ratio | Distance (moisture availability) | 0.08 | <0.001 | 0.13 |
| Bhattacharyya distance | Distance (moisture availability) | 0.26 | <0.001 | 0.18 |

We measured the pairwise functional turnover between any two sites as the Bhattacharyya distance between their MVN hypervolumes, and according to Equation 7, partitioned it into the Mahalanobis distance (which measures the standardized distance between community means) and the determinant ratio (which measures the differences in functional alpha diversity) (see a two-community example in Fig. 2b). The ratio of Mahalanobis distance to determinant ratio (MD/DR) reflects the relative importance of difference in community means and trait diversity differences in driving the overall community-level functional turnover. The median ratio MD/DR across all pairs of sites is 0.63, suggesting functional turnover among communities is mainly driven by the variation in functional alpha-diversity. The functional turnover is more determined by moisture availability than by minimum temperature (Fig. 2e, f). Multiple regression on distance matrices suggests that the Mahalanobis distance is correlated with both minimum temperature and moisture availability, but the determinant ratio is only correlated with moisture availability (Table 1).

### Species level: environmental niches

We used the presented methodology to analyze and compare the three-dimensional environmental niches (temperature, precipitation, and EVI) of 260 bird species in the New World. We partitioned the total environmental niche volume for each single species into the three environmental variable variance components and the correlation component. For a two-species example, we attributed the difference in niche volume between the white-throated swift and the tricolored blackbird to differences in temperature variance, precipitation variance, EVI variance and the correlation component (Fig. 3a). For the 260-species, assessment, we found that the coefficient of variation for each of the niche volume components among 260 bird species was 0.79 for precipitation variance, 0.67 for temperature variance, 0.44 for EVI variance, and 0.22 for the correlation component, suggesting that precipitation niche breadth variation contributes most to the species-level niche volume variation. Range size and centroid latitude are the two main drivers of realized environmental niche volume (Fig. 3c, d), but the predictors for each niche volume component are different: the precipitation variance and the correlation component are only correlated with the centroid latitude while the temperature variance and the EVI variance are correlated with both centroid latitude and range size (Table 2). The median ratio between Mahalanobis distance and determinant ratio is 2.43, suggesting that the distance between niche centroids is more important than niche volume differences in driving overall environmental niche dissimilarity among the 260 bird species. When the multiple regression on distance matrices (MRM) is applied, different components of the Bhattacharyya distance have different determining factors: the Mahalanobis distance is only correlated with centroid latitude; while the determinant ratio is correlated with both range size and centroid latitude (Table 2).

**Figure 3.**
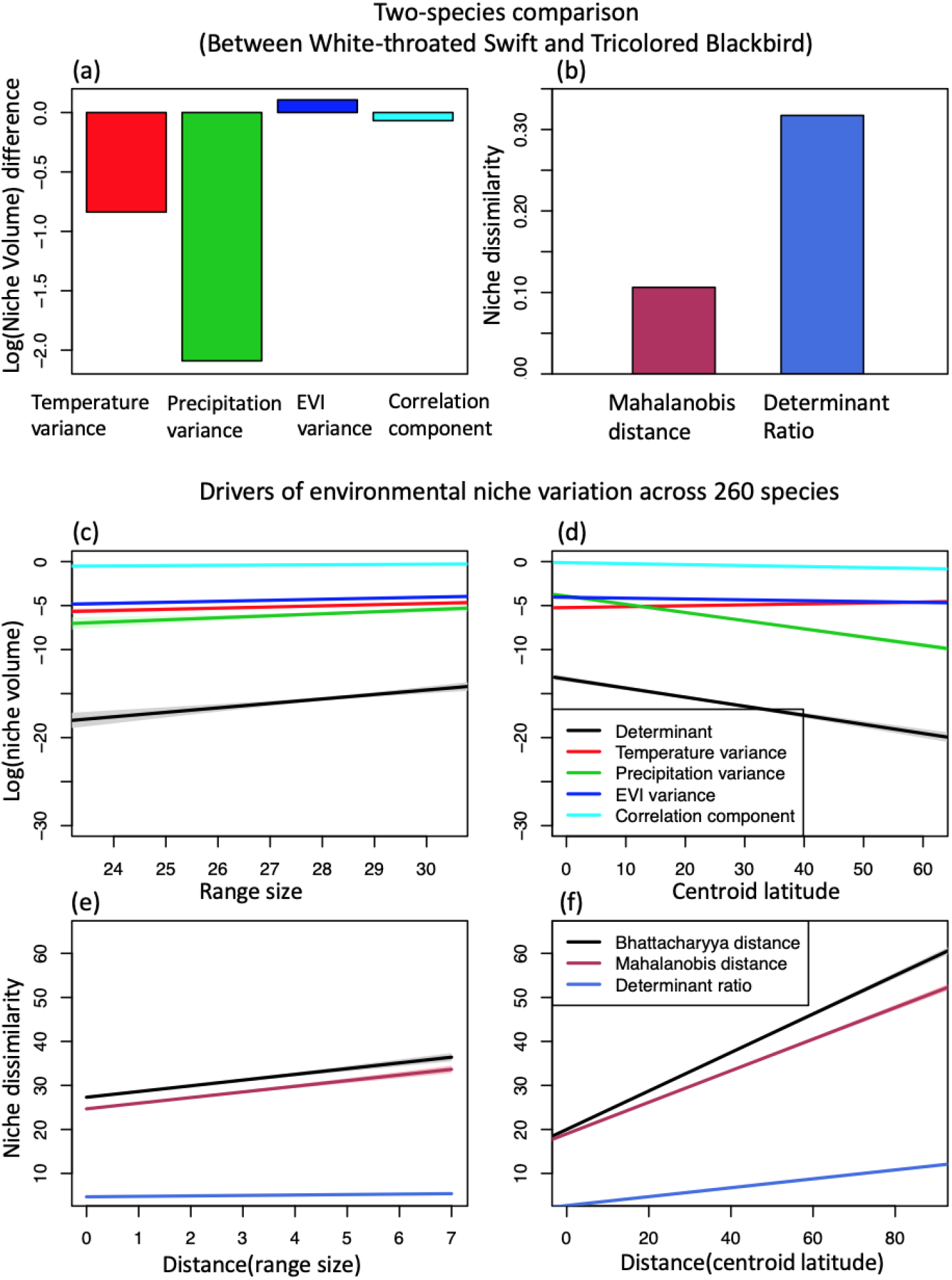
Species-level environmental niche assessment. (a) Component-wise environmental niche volume differences and (b) Bhattacharyya distance of environmental niche dissimilarity between two example species (White-throated Swift and Tricolored Blackbird) (c)-(f) Change of niche volume and pairwise niche dissimilarity among 260 bird species with range size and centroid latitude. Solid lines are the fitted linear regression. Shaded area represents 95% confidence interval of linear regression. See table 2 for summary statistics.

**Table 2.** Summary statistics for multiple linear regression of environmental niche volumes, and Multiple regression on distance matrices on environmental niche dissimilarity. Area, temperature, precipitation, EVI variances, the correlation component and the determinant are log-transformed. Adjusted R^2^ is used. Only significant predictors are reported.

| response | predictor | estimate | p-value | $R^2$ |
| --- | --- | --- | --- | --- |
| Temperature variance | Area | 1.13 | <0.001 | 0.14 |
|  | Centroid latitude | 0.84 | <0.001 |  |
| Precipitation variance | Centroid latitude | -5.72 | <0.001 | 0.69 |
| EVI variance | Area | 0.70 | <0.001 | 0.13 |
|  | Centroid latitude | -0.46 | <0.001 |  |
| Correlation component | Centroid latitude | -0.67 | <0.001 | 0.22 |
| Determinant | Area | 2.09 | <0.001 | 0.55 |
|  | Centroid latitude | -5.97 | <0.001 |  |
| Mahalanobis distance | Distance (Centroid latitude) | 32.62 | <0.001 | 0.13 |
| Determinant ratio | Distance (Area) | -2.66 | <0.001 | 0.23 |
|  | Distance (Centroid latitude) | 9.78 | <0.001 |  |
| Bhattacharyya distance | Distance (Centroid latitude) | 40.36 | <0.001 | 0.15 |

### Comparison with other metrics

Community-level functional traits: The determinant of trait covariance matrix as the measure of functional diversity had the highest correlation with the area of the 95% isopleth of a kernel density derived hypervolume (0.89), followed by functional dispersion (0.79) and minimum convex hull (0.65) (additional methods in Fig. 4a). The Bhattacharyya distance had a similar correlation with Euclidean centroid distance (0.70), Jaccard similarity (−0.69) and Sorensen similarity (−0.71). The Mahalanobis distance component was more correlated with Euclidean centroid distance (0.73) than with Jaccard similarity (−0.57) or Sorensen similarity (−0.59), while the determinant ratio component had a relatively low correlation with the Euclidean centroid distance (0.42; Fig. 4b). All components were uncorrelated with minimum distance between hypervolumes (Fig. 4b).

**Figure 4.**
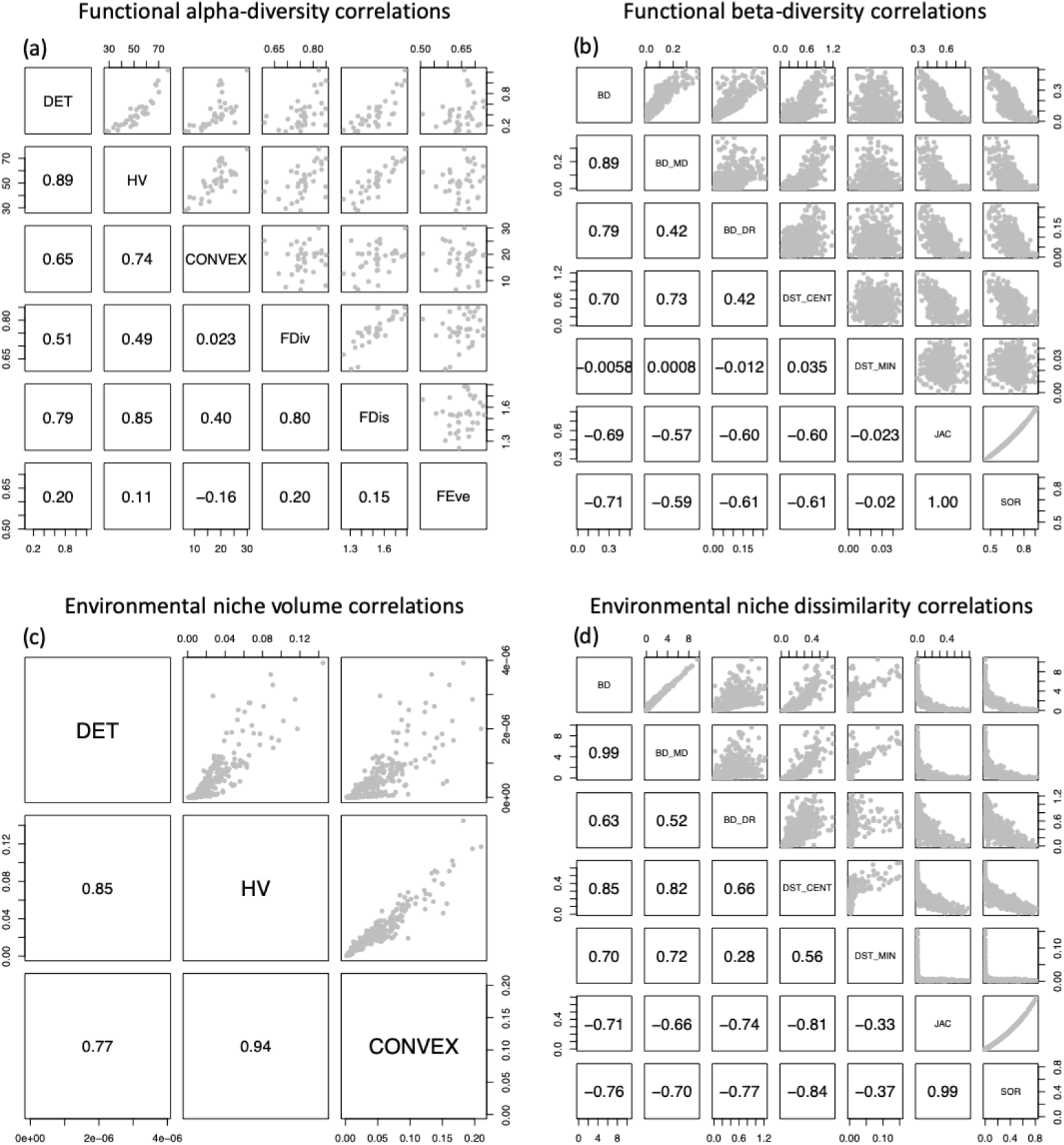
(a) Correlations among functional alpha-diversity measures for the 36 sites. (b) Correlations among pairwise functional turnover measures. (c) Correlations among environmental niche volume measures for the 260 bird species. (d) Correlations among pairwise environmental niche dissimilarity measures. ‘DET’ stands for the determinant of the environmental niche or functional trait covariance matrix, ‘HV’ for kernel density derived hypervolume, ‘CONVEX’ for minimum convex hull, ‘FDiv’ for functional divergence, ‘FDis’ for functional dispersion, ‘FEve’ for functional evenness. ‘BD’ stands for Bhattacharyya distance, ‘BD_MD’ for the Mahalanobis distance component of BD, ‘BD_DR” for the determinant ratio component of BD, ‘DST_CENT’ for Euclidean distance between centroids, ‘DST_MIN’ for minimum distance between kernel density derived hypervolumes, ‘JAC’ for Jaccard similarity, ‘SOR’ for Sorensen similarity.

Species-level environmental niche: the determinant of the environmental niche covariance matrix correlates most strongly with the area of the 95% isopleth of the kernel density derived hypervolume (0.85), followed by the minimum convex hull (0.77, Fig. 4c). The Bhattacharyya distance and its Mahalanobis distance component are most correlated with the Euclidean centroid distance (0.85 and 0.82 respectively); the determinant ratio is similarly correlated with most alternative metrics except for minimum distance between hypervolumes (Fig. 4d).

## Discussion

We proposed a new parametric hypervolume framework based on multivariate normal distributions and corresponding size and dissimilarity metrics. The greatest advantage of the MVN hypervolume approach is the analytical clarity and insights provided by the partitioned components of the niche size and dissimilarity metrics. The MVNH framework is also significantly faster than the widely used KDE approach. For example, using the ‘Iris’ dataset in R (a dataset with 150 rows and 3 columns), it takes 6 seconds to compute the hypervolume using KDE but only 0.001 seconds to compute the hypervolume using MVNH. Although the determinant of the covariance matrix has been used as a total niche volume measure in some studies (Soberon & Nakamura, 2009; La Sorte et al., 2018), it has never been illustrated that the determinant could be interpreted as the product of individual niche variances and a correlation component. The Bhattacharyya distance has been used as a measure of home range overlap in animal movements (Fieberg & Kochanny, 2005; Winner et al., 2018). We suggest that its use in trait or environmental space (Lu et al., 1989; Minami & Shimizu, 1999), and in particular the interpretation of the partitioned components, offers valuable biological insights. Specifically, the Mahalanobis distance between hypervolume centroids can be interpreted as the standardized distance between optimal traits of two communities or between the optimal environmental conditions of two species under the assumptions that mean community traits represents the optimal traits under equilibrium (Blonder et al., 2015) and that niche centroid has the highest fitness for a species (Sagarin, Gaines, & Gaylord, 2006). The determinant ratio can be interpreted as the functional diversity difference between communities or as niche volume differences between species. The Bhattacharyya distance (BD) not only avoids choosing ad hoc algorithm parameters, but more importantly reconciles the limitations of the two commonly used classes of dissimilarity metrics. When Jaccard or Sorensen indices become uninformative for disjunct hypervolumes, the BD incorporates the distance information through the Mahalanobis distance component. When the centroid distance metric becomes uninformative for two close and overlapping hypervolumes, the BD incorporates the shape information through the determinant ratio component.

The good concordance between our method and the kernel density method across each of our studies suggests that the determinant as a measure of niche volume is robust to different distributions of data. For niche dissimilarity measures, the moderate amount of correlation between the partitioned components of BD with the Jaccard, Sorensen and the centroid Euclidean distances shows that the partitioned components of BD both capture a part of the shape information of the hypervolume. Therefore, our BD metrics combine the important interpretations of both centroid Euclidean distance metrics and the Jaccard and Sorensen metrics of niche dissimilarity. More importantly, the relative importance of centroid difference and volume difference are directly comparable using the BD metrics.

Other metrics such as the MacArthur-Levins measure (Equation 9), the Pianka’s measure (Equation 10) and the Morisita’s measure (Equation 11) could be similarly partitioned into a Mahalanobis distance component and a determinant ratio component (Lu et al., 1989), though their properties are less well known than the Bhattacharyya distance (Minami & Shimizu, 1999; Winner et al., 2018). The shared Mahalanobis distance component in all these metrics could be further partitioned into standardized Euclidean distance along different PCA axes according to a previously proposed partitioning framework (Calenge et al., 2008; Mahony et al., 2017), which measures the relative importance of each PCA axis in determining the overall Mahalanobis distance.

### Case studies

Through the partitioning framework, our methods provide a novel understanding of each component driver of community functional diversity or species environmental niche volume. In the functional diversity example we demonstrated that for the 36 annual plant communities the constraints on combination of traits is the major driver of functional alpha diversity variation rather than the change of individual trait variances, which in turn is shaped by moisture availability; the major driver of functional community turnover among is the variation of functional alpha-diversity rather than the mean trait differences between communities. In the species environmental niche example, our method shows that the major driver of niche volume variation is the precipitation niche breadth variation (Fig. 3d) which is determined by centroid latitude of species’ range. We also show that the range size-niche breadth relationship of total environmental niche breadth is mainly driven by the range size-temperature niche breadth and range size-EVI niche breadth relationship (Table 2).

### Caveats

A common limitation of all hypervolume metrics is the curse of dimensionality and collinearity among axes (Blonder et al., 2014; D’Andrea & Ostling, 2016). The use of the determinant is especially sensitive to collinearity because any pair of fully dependent columns or rows in a covariance matrix will necessarily result in a determinant of zero. Increasing dimensions of trait or environmental data will increase the chance of collinearity and is also more likely to violate the multivariate normal assumption. Such an issue could be resolved by choosing the biologically relevant variables a priori or using common dimension reduction techniques applied in non-parametric methods such as PCA (Junker et al., 2016). Because PCA creates orthogonal axes, the correlation component will always equal 1 if PCA is applied, and thus it is only recommended when the correlation among axes is of no interest to the study.

### Potential

The presented size and dissimilarity measures are to our knowledge the first parametric metrics that allow a partitioning of n-dimensional hypervolumes. The partitioning of total niche volume and niche dissimilarity enables more rigorous testing of ecological and evolutionary hypotheses beyond those we explored in our two case studies. For example, when investigating the spatial scaling of niche volume (Pearman, Guisan, Broennimann, & Randin, 2008; Connor et al., 2018), the scaling relationship could be broken down to the scaling of univariate niche variances and correlation component, each of which could be linked to different sets of predictors. When studying niche shifts during biotic invasion (Lauzeral et al., 2011; Hill, Hoffmann, Macfadyen, Umina, & Elith, 2012; Wiens, Litvinenko, Harris, & Jezkova, 2019), the BD could be used to determine whether the niche shift is mainly caused by shifts in niche centroids or caused by volume expansion or reduction; the determinant could be used to assess which niche axis is most responsible for the total volume expansion or reduction. In niche evolution studies, the evolution of niche dissimilarity (Nunes & Pearson, 2017; Castro-Insua et al., 2018) between two species could be traced to contributions by niche breadth or center differences to the total niche divergence. The parametric measures also have closer links to theory as theoretical predictions of niche evolution are usually derived from covariance matrixes with multivariate normal approximations (Lande & Arnold, 1983; Caetano & Harmon, 2019). In other words, the newly proposed metrics have the potential to provide more mechanistic understanding of biodiversity patterns and niche evolution, and to pave the way for more accurate forecasts for biodiversity change.

## Acknowledgement

We thank Jetz Lab members for discussion of the work, especially Richard Li, Aurore Maureaud and Ben Carlson for comments on the first draft.

## Authors’ contributions

ML and WJ conceived the ideas. ML analyzed the data with substantial inputs from KW and WJ. ML led the writing of the manuscript. All authors contributed critically to the drafts and gave final approval for publication.

